# Host DNA repair factors empower a mechanism of antiviral nucleoside analog resistance

**DOI:** 10.64898/2026.05.21.726832

**Authors:** Pierce Longmire, Han Chen, David R. McKinzey, Mamata Savanagouder, Noelle N. Kosarek, Jean M. Pesola, Carly A. Bobak, Giovanni Bosco, Felicia Goodrum, Donald M. Coen

## Abstract

How host functions affect resistance to antiviral drugs is poorly understood. Ganciclovir, a chain-terminating nucleoside analog, is a first-line therapy against human cytomegalovirus, a widespread herpesvirus that causes life-threatening disease in immunocompromised individuals and newborns. Ganciclovir resistance, which is caused by mutations that affect the viral kinase, UL97 and/or the viral polymerase, UL54, can cause treatment failures. Among these mutations, those reducing the exonuclease activity of the viral DNA polymerase permit ganciclovir incorporation without chain termination. However, the fate of DNA strands containing the incorporated nucleotide analog is unknown. We show here that template DNA containing ganciclovir fails to support DNA synthesis of the complementary strand by exonuclease-mutant polymerase. Moreover, while DNA synthesis and ganciclovir incorporation are limited in drug-treated fibroblasts infected by virus with wild-type polymerase, an exonuclease-resistant mutant virus can better synthesize full-length genomes and incorporate substantially more ganciclovir into DNA. Notably, ganciclovir is lost from DNA when drug is removed, suggesting that ganciclovir-containing templates are repaired. We identify the host nucleotide excision repair component, XPA, and the repair enzyme, polymerase kappa, as each being necessary for mutant virus ganciclovir resistance and polymerase kappa as being required for the mutant’s cidofovir resistance, demonstrating a role for host DNA repair machinery in a mechanism of antiviral resistance. We propose a model for this mechanism, which has relevance for at least one other antiviral drug and likely other nucleoside analog therapeutics, and highlights the participation of host DNA repair machinery during human cytomegalovirus DNA replication.

**IMPORTANCE:** Nucleoside analogues such as ganciclovir, which is a leading drug for preventing and treating human cytomegalovirus, are a critical defense against viral diseases, but antiviral resistance often results in treatment failures. This study reveals a critical role for host DNA repair in a mechanism of resistance to ganciclovir, and identifies at least one specific repair pathway that permits viral DNA synthesis in the presence of ganciclovir, defining a mechanism by which cellular DNA repair pathways conspire to enable antiviral drug resistance. This mechanism is relevant to at least one other antiviral drug and may apply to other antiviral and anticancer agents. The study also showcases the participation of host DNA repair machinery during human cytomegalovirus DNA synthesis.

## INTRODUCTION

Investigating biological processes is invaluable for the discovery of therapeutics and for understanding their mechanisms of action and resistance. Similarly, studies of therapeutics can be invaluable for understanding biological mechanisms. A biological process shared by all free-living organisms is DNA repair. Notably, many viruses also are subject to DNA repair and/or make use of cellular repair machinery during their life cycles. Human cytomegalovirus (HCMV) is a herpesvirus with a DNA genome of 230-240 kilobase pairs (kbp) that causes a variety of severe diseases in immunologically naïve or immunocompromised individuals(1). HCMV encodes numerous proteins that participate in viral DNA synthesis, including a DNA polymerase that consists of a catalytic subunit, UL54 (Pol) with both 5’-3’ polymerase and 3’-5’ exonuclease activities, and an accessory subunit, UL44, that promotes long chain DNA synthesis(2–5). The virus also encodes a DNA repair enzyme, uracil DNA glycosylase, that is important for removal of uracil from DNA and for viral DNA synthesis(6–8). Despite the presence of this viral machinery, a number of studies have indicated that HCMV recruits host DNA replication and repair factors, including proteins involved in homologous recombination repair (HR), nucleotide excision repair (NER), and translesion synthesis (TLS), to the sites of viral DNA synthesis in the host cell nucleus to regulate viral DNA synthesis and the production of infectious virus(9–15). Moreover, some host factors, including TLS DNA polymerases(9, 15), contribute to replication and repair of viral genomes in difficult-to-replicate regions.

First-line therapies for HCMV include ganciclovir (GCV) and its more orally bioavailable prodrug, valganciclovir (reviewed in (16)). Like many widely used antiviral, antifungal, and anticancer agents(17), GCV is a nucleoside analog. GCV mimics deoxyguanosine but in place of deoxyribose contains an acyclic sugar moiety lacking the equivalent of a 2’-position but having the equivalent of a 3’-hydroxyl group(18). In HCMV-infected cells, GCV is phosphorylated to its monophosphate (GCV-MP) by the virus-encoded UL97 kinase, and subsequently to GCV-diphosphate and GCV-triphosphate (GCV-TP) by cellular kinases (reviewed in (16)). Consistent with the production of short chains of viral DNA in infected cells(19, 20), biochemical assays have shown that GCV-TP competes with deoxyguanosine triphosphate for incorporation into viral DNA by Pol, causing chain termination, but only after incorporation of the subsequent (N+1) nucleotide(18, 21–23). Further biochemical studies indicate that after incorporating GCV and the N+1 nucleotide, the exonuclease activity of Pol is unable to degrade the primer strand(24). Subsequent polymerase activity greatly slows, while the exonuclease activity rapidly removes the incorporated N+2 nucleotide and generates its monophosphate (idling)(24, 25). The net result is termination at the N+1 position. However, mutant Pols with insufficient exonuclease activity (Exo-mutants) overcome chain termination and synthesize full length products(24, 25). The majority of GCV-resistant viruses with *pol* mutations have alterations in the exonuclease domain of the polymerase(26).

These findings imply that GCV-resistant (GCV^r^), Exo-mutant viruses would internally incorporate GCV during synthesis of full-length genomes. Based on NMR structural analysis of oligonucleotides containing internally incorporated GCV(27), this is expected to distort the DNA backbone with deleterious consequences to replication of the genome and gene expression. Yet, resistance occurs, suggesting that the viral Pol can copy GCV-containing template DNA or that there are DNA repair mechanisms that remove the GCV and replace it with normal deoxynucleotides.

Evidence that host cells have the capacity to repair GCV-containing genomes comes from studies of GCV as an anti-cancer therapy in combination with transduction of tumor cells with herpes simplex virus 1 thymidine kinase, which efficiently phosphorylates GCV (GCV-HSV-TK)(28)(reviewed in (29)). Several lines of evidence indicate that DNA repair mechanisms reduce cytotoxicity of GCV-HSV-TK(30–33), limiting its anti-cancer potential. Notably, in these studies, although host DNA polymerases also terminate chain elongation after incorporation of GCV and the N+1 nucleotide in vitro(18, 21, 23), GCV was nevertheless incorporated into high molecular weight DNA, and treated cells completed one cell division(34, 35). GCV-induced cytotoxicity and cell death are manifested during the following cell cycle(34, 36, 37). Incorporation of GCV into templates for subsequent DNA synthesis is a critical event in DNA damage and cell death, which might be initiated by repair processes on those templates(34, 38).

We hypothesized that incorporation of GCV could be deleterious to Exo-mutant viruses and would thus need to be repaired to permit their ganciclovir-resistance. To explore these hypotheses, we undertook biochemical experiments to test the ability of HCMV Pol to copy templates with incorporated GCV, and radiolabeling studies to determine if incorporated GCV is removed from Exo-mutant genomes during infection. Finally, we utilized somatic cell genetics and RNA interference studies to identify host DNA repair factors important for repairing viral genomes and enabling GCV-resistance of an Exo-mutant virus. *Prior to this study, the textbook view of GCV-resistance was entirely attributed to mutations in the viral polymerase (UL54) or the viral kinase (UL97). We have identified both a host pathway and two host factors within the pathway that contribute to a mechanism of antiviral resistance*.

## RESULTS

### GCV-containing DNA template strands block DNA synthesis catalyzed by HCMV Pol

We previously showed that WT Pol extends DNA primer strands containing GCV by one or two additional nucleotides so slowly that idling, due to more rapid exonuclease activity, leads to a net result of termination at the N+1 position(24, 25). By contrast, Exo-mutant Pols corresponding to those encoded by GCV^r^ mutant viruses, efficiently elongate DNA primer strands containing GCV, which could result in GCV remaining in the template strand for the next round of DNA synthesis (24). Therefore, we asked if GCV in the DNA template strand would impede DNA synthesis *in vitro*. For this purpose, we used purified WT Pol and two previously described Exo-mutant Pols (F412V and L545S)(24), all expressed as glutathione-S-transferase (GST) fusion proteins. We also used a previously described GST-tagged mutant Pol that, rather than being an Exo-mutant, contains a GCV^r^ substitution in the polymerase thumb domain (A987G) and extends GCV-containing DNA primers faster than WT Pol, thus overcoming GCV-induced chain termination to allow efficient DNA chain elongation by a mechanism other than loss of exonuclease activity(25). As a control we used exonuclease-deficient bacteriophage T7 DNA polymerase as it has been reported to be able to bypass certain adducts in the template strand during DNA synthesis(39, 40).

We synthesized a 39-mer hairpin primer-template (T5), in which GCV is the first unpaired nucleotide of the template, 3’ to the double stranded primer region (Fig.1). As a control to determine whether the various polymerase preparations were comparably active, we also used the T1 template(24), which has no GCV residues. Using either the T1 or T5 primer-template radiolabeled on its 5’-end, in the presence of all four deoxynucleoside triphosphates (dNTPs), the control bacteriophage T7 DNA polymerase elongated both the T1 and T5 primer-templates efficiently, resulting in full-length products (∼85% and ∼100%, respectively, from densitometry) along with some minor, smaller extension products (lanes 2 and 4). Minor species shorter than the primer-templates are also evident that likely represent contaminants that arose during oligonucleotide synthesis. These are also seen in lanes 3 and 10 that contain the T5 primer-template alone. The ability of the T7 enzyme to synthesize DNA across from GCV is consistent with its ability to bypass other adducts on the template strand (39, 40). WT Pol and all three mutant HCMV Pols (which were tested in the presence of the presumptive HCMV processivity factor, UL44) also efficiently elongated the T1 primer-template lacking GCV (lanes 6-9) indicated by the ∼100% full-length or longer products. The longer species were detected previously in reactions using Exo mutants (24) and might arise due to non-templated addition of a nucleotide, which occurs with other exonuclease-deficient polymerases(41), or possibly due to slippage of the enzymes on the run of six dT residues at the end of the template. Thus, the enzymes were active. None of the HCMV Pols efficiently elongated the T5 primer-template (lanes 10-14). Less than 1% full-length products were observed for WT, F412V, L545S Pols and 31% for the A987G Pol. However, some minor products, suggestive of dNTP incorporation opposite the GCV in the template strand, formed in reactions with the mutant polymerases. Even then, ∼80% of the DNA primers were not extended.

**Figure 1.**
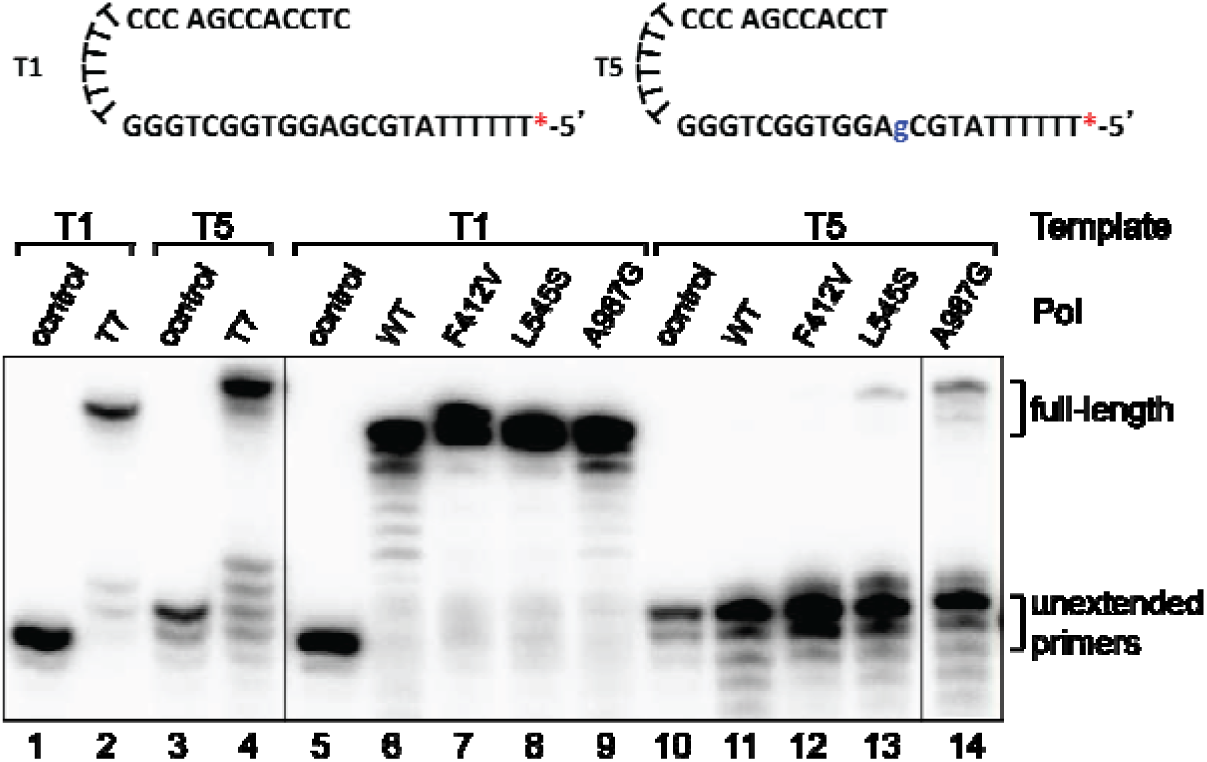
Template-incorporated GCV reduces DNA extension by WT and Exo-mutant HCMV Pols. Radiolabeled primer templates T1 or T5 (top; the position of the radiolabel in both T1 and T5 is shown with a red asterisk and the position of GCV in T5 is indicated by a lower case blue g) were incubated with dNTPs and each of the indicated WT or Exo-mutant HCMV Pols (lanes 6-9 with T1; lanes 11-14 with T5) in the presence of UL44, or as a control, with dNTPs and exonuclease-deficient T7 DNA polymerase (lane 2 with T1 and lane 4 with T5) at 37° C for 10 min, and the products were analyzed alongside untreated T1 (lanes 1 and 5) and T5 (lanes 3 and 10) using polyacrylamide gel electrophoresis and autoradiography. The Pol and primer template used are indicated at the top of each lane. The positions of unextended primers and full-length products are indicated on the right. The vertical lines between lanes 4 and 5 and between lanes 13 and 14 indicate a lane deleted from the figure that was a duplicate of lane 4 and a lane that was not used because of a bubble in the gel, respectively.

Taken together, these data show that neither WT nor Exo-mutant Pols efficiently synthesized DNA from a primer-template that contains GCV in the template strand, suggesting that GCV incorporation into template DNA results in a lesion that could block subsequent viral DNA synthesis.

### GCV incorporated into viral DNA during Exo-mutant virus infection is subsequently removed

Given the results above, we then investigated the fate of incorporated GCV in cells infected with Exo-mutant virus F412V. For this purpose, we engineered the F412V substitution into a GFP-expressing version of the AD169 strain of HCMV using bacterial artificial chromosome (BAC) technology and then engineered a rescued derivative of F412V, F412V_R, from the F412V BAC. The F412V mutant polymerase failed to synthesize DNA across from GCV incorporated in the template strand (Fig. 1) and has previously been shown to lack detectable exonuclease activity and to readily extend GCV-containing primers in biochemical assays(24). Moreover, the F412V substitution has been found in a clinical isolate and has been reported to confer ∼4-fold resistance in antiviral assays (42, 43), which is intermediate and within ∼2-fold of fold-resistance values (1.9- to 8.0-fold) conferred by UL54 mutations in clinical isolates (26).

The BAC-derived F412V mutant virus exhibited a nearly five-fold higher ED_50_ to GCV compared with its WT parent and F412V_R using yield reduction assays in human foreskin fibroblasts (Fig. S1A). These differences were significant and consistent with the results of previous plaque reduction assays of other F412V HCMV mutants in human fibroblasts(42, 43). Of note, in this experiment and in the experiments shown in Figs. 2, S2, and 4 below, GCV was administered to cells in medium containing 1% DMSO and the half maximal effective dose (ED_50_) values for WT were ∼0.1 μM. In other experiments (Figs. 3 and 5-6, shown below), the DMSO concentration used was 0.1% and the ED_50_ values for WT were ∼1 μM. In a side-by-side comparison, 1% DMSO was found to decrease the ED_50_ for WT virus (Fig. S1B-C) during infection of MRC-5 primary human lung fibroblasts relative to 0.1% DMSO, explaining much of the difference in ED_50_ values. To determine if GCV-containing viral DNA could be repaired, we sought to observe both GCV incorporation into and removal from the F412V genome. To avoid either affecting viral DNA synthesis so severely or so little as to make GCV incorporation and removal undetectable, we tested two doses of GCV, 0.3 μM and 0.6 μM—that were either just below or just above the ED_50_ of GCV for the F412V mutant yet greater than the ED_50_ of GCV for WT—in our assay (Fig. S1A) for effects on the kinetics of viral replication at a multiplicity of infection (MOI) of 2. In these pilot experiments, in the absence of drug, F412V exhibited replication kinetics similar to that of WT (Fig. S2A, B). We found that treatment with 0.3 μM GCV led to only modest effects on both WT (∼3-fold) and mutant (∼1.4-fold) yields at 96 hours post-infection (hpi) (Fig. S2A). On the other hand, 0.6 μM GCV led to a 20-fold reduction in WT yield at 96 hpi while reducing the viral yield of F412V by only ∼3-fold (Fig. S2B). We chose this concentration for radiolabeling infected cell DNA with [^3^H]-GCV.

**Figure 2.**
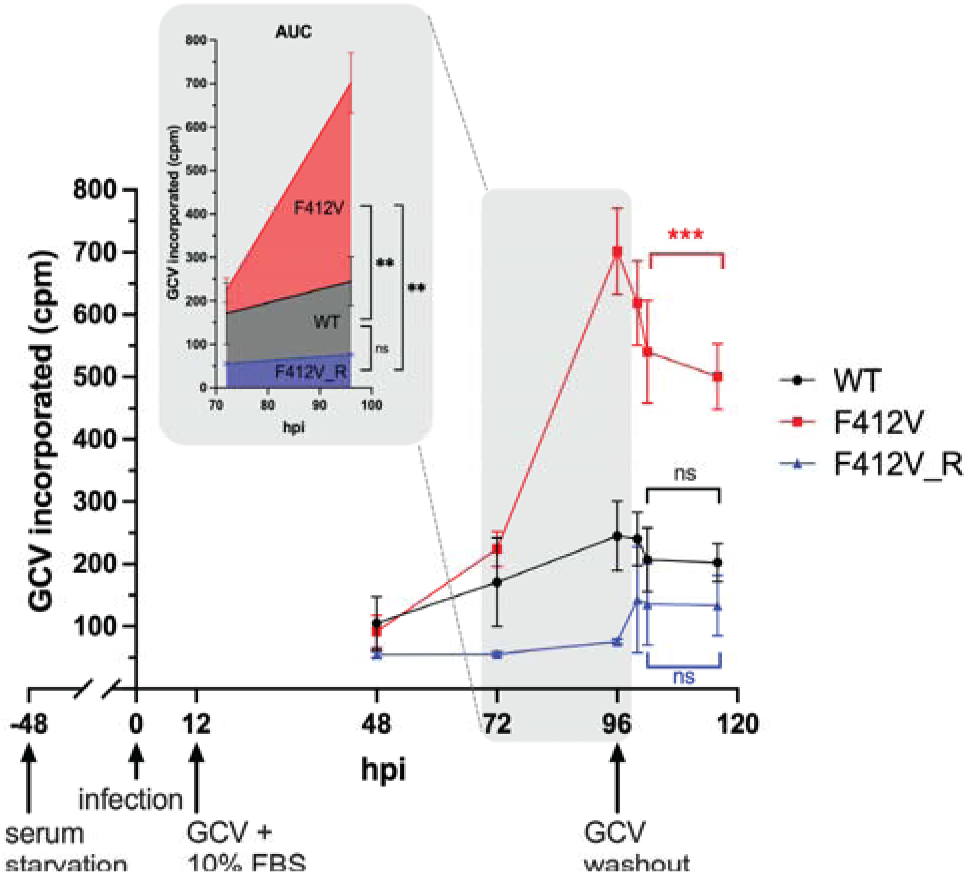
Loss of incorporated GCV from replicating Exo-mutant HCMV DNA. Human foreskin fibroblast cells, serum-starved for 48 h, were infected with the indicated viruses at an MOI of 4 for two hours, and then low serum medium was added back. At 12 hpi ^3^H radiolabeled GCV (0.6 μM) was added in medium containing 10% FBS, and at 96 hpi medium was removed, cells rinsed, and medium without GCV was added. Individual plates of cells were harvested at the indicated time points, and the amounts of radiolabeled GCV in counts per minute (cpm) incorporated into DNA per plate were measured. Background cpm “incorporated” in the absence of radiolabel (25 cpm) were subtracted, and there were ∼75 background-subtracted cpm incorporated into DNA per plate in mock infected cells at 96 hpi. Data are from triplicate experiments. Error bars represent standard deviations (S.D.). As depicted in the inset graph, areas under the curves (AUCs) for GCV incorporation were calculated for each virus between 72 and 96 hpi. These areas were analyzed via one-way ANOVA, allowing for unequal S.D., and followed by Dunnett’s T3 multiple comparison tests. Repeated measures two-way ANOVA (matched by experiment) followed by Šídák’s multiple comparison tests were used to evaluate the degree of GCV washout at 116 hpi relative to 96 hpi (***, p<0.001; **, p<0.01; ns, not significant).

**Figure 3.**
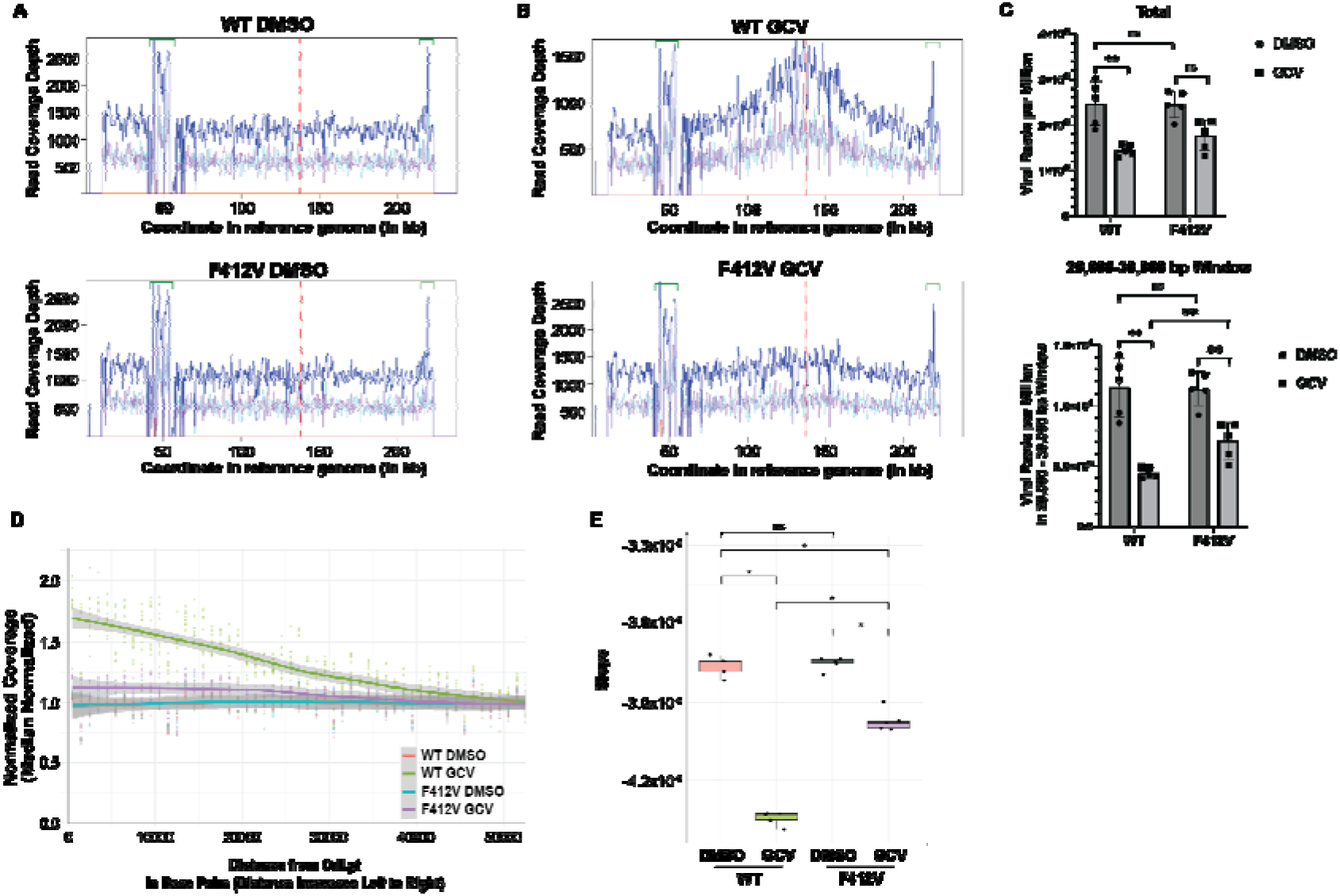
GCV incorporation terminates vDNA synthesis around the oriLyt in WT infection, but not F412V Exo-mutant virus infection. Primary lung fibroblasts (MRC-5) were infected with WT or F412V at an MOI of 1. At 2 hpi, virus was washed out and media were added containing either 0.1% DMSO or 0.1% DMSO with 1.6 µM GCV. At 96 hpi, cells were harvested and total DNA was isolated for Illumina sequencing. After sequence alignment and removal of reads mapping to the human genome, viral reads were quantified and plotted for each condition. (A and B) Representative sequence coverage plots of total unique (dark blue) viral reads from WT (A) and F412V (B) infected cells treated with DMSO vehicle (top) or GCV in vehicle (bottom). The red dotted line represents the location of the *oriLyt* (location ∼140,000bp in the genome). Top (cyan) and bottom (purple) strand reads are depicted separately. Only unique reads that map to only one location in the reference genome were counted. (C) Total reads per million (top) and reads distant from *oriLyt* (bottom) were quantified. Non-parametric Wilcoxon testing was used to determine statistical significance. The resulting p-values were adjusted using the Benjamini-Hochberg method: **, p<0.01; ns, not significant) (D) Sequence coverage decay plots representing the normalized coverage at the *oriLyt* location up to 50kb moving away from the *oriLyt* in both directions. (E) Distribution of decay slopes and pairwise comparisons of adjusted p-values of WT and F412V treated with either DMSO or GCV. The slopes estimated per sample were compared across groups using pairwise permutation testing (with BH correction) to assess significant differences in replication bias patterns between conditions: *, p<0.05; ns, not significant.

**Figure 4.**
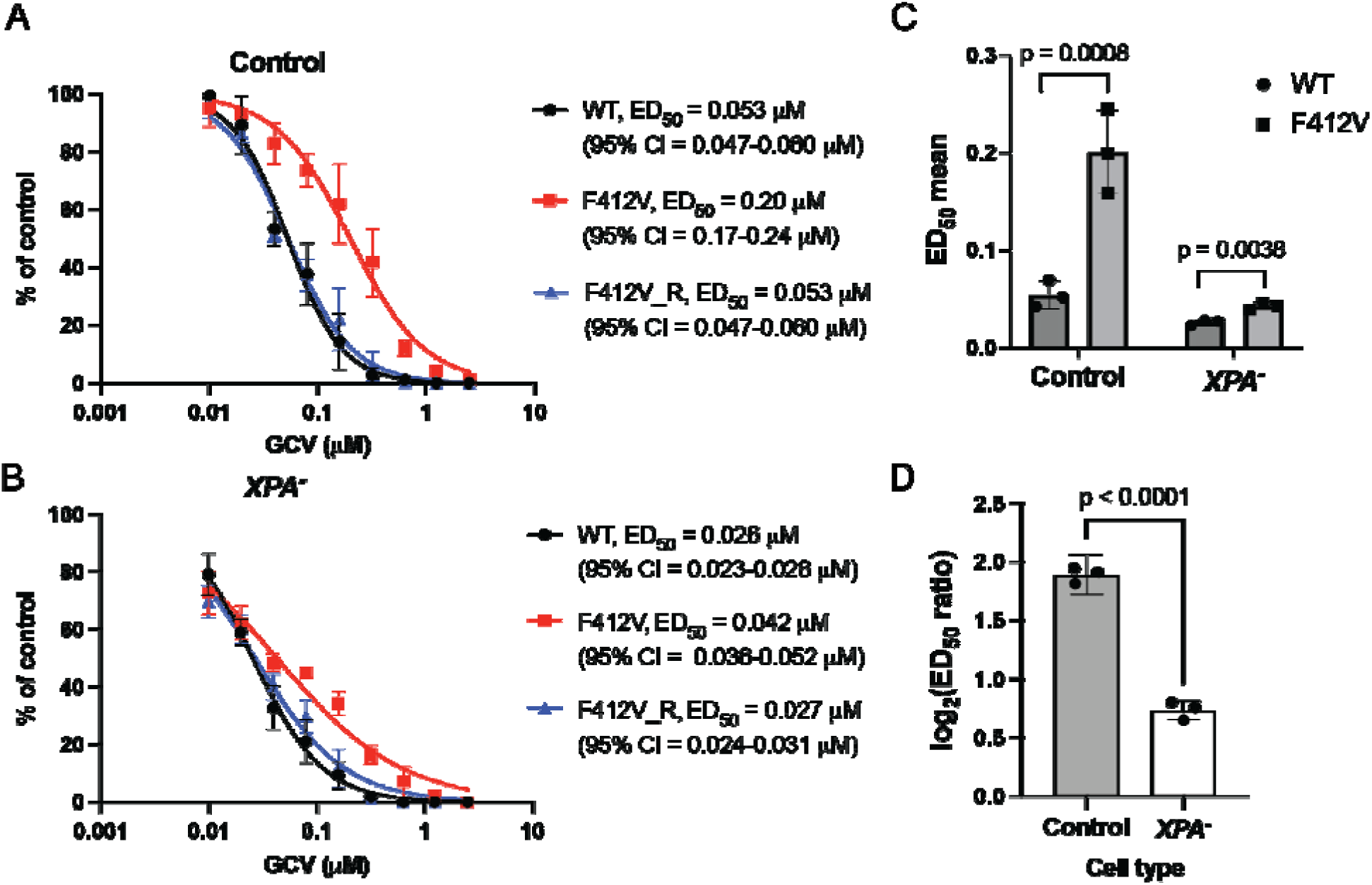
Reduced resistance to GCV of Exo-mutant HCMV in XPA deficient cells. (A-B) Control cells (A) and *XPA^-^* cells (B) were infected at MOI 2 with WT HCMV, the Exo-mutant virus F412V or its rescued virus F412V_R, respectively. After 2 h incubations, the viruses were removed and media containing various concentrations of GCV were added. At 5 dpi, the supernatants were harvested. The viruses were titrated on HFF cells and the yield of virus at each concentration of GCV was calculated using standard plaque assay. The ED_50_ values for GCV inhibition and their 95% confidence intervals (CI) presented in these two panels were determined from triplicate experiments by least squares regression with variable slope. (C) Curves were similarly fitted for WT and F412V in each independent experiment to obtain three ED_50_ values for each cell type. Those ED_50_ values from three biological replicate experiments were used for a ratio paired t test comparing viruses within each cell type and the Holm-Šídák method was used to correct for multiple comparisons. ED_50_ mean values are plotted with standard deviation error bars. (D) From each experiment, for each cell type, the ratio of the ED_50_ for WT virus to the ED_50_ for F412V was calculated. Because ED_50_ ratios typically do not follow a normal distribution, which is necessary for t tests, while logarithms of such values typically do follow a normal distribution(74), we plotted log_2_ values and used them for an unpaired t test with Welch’s correction. Mean log_2_(ED_50_ ratio) values are shown with 95% CI error bars.

**Figure 5.**
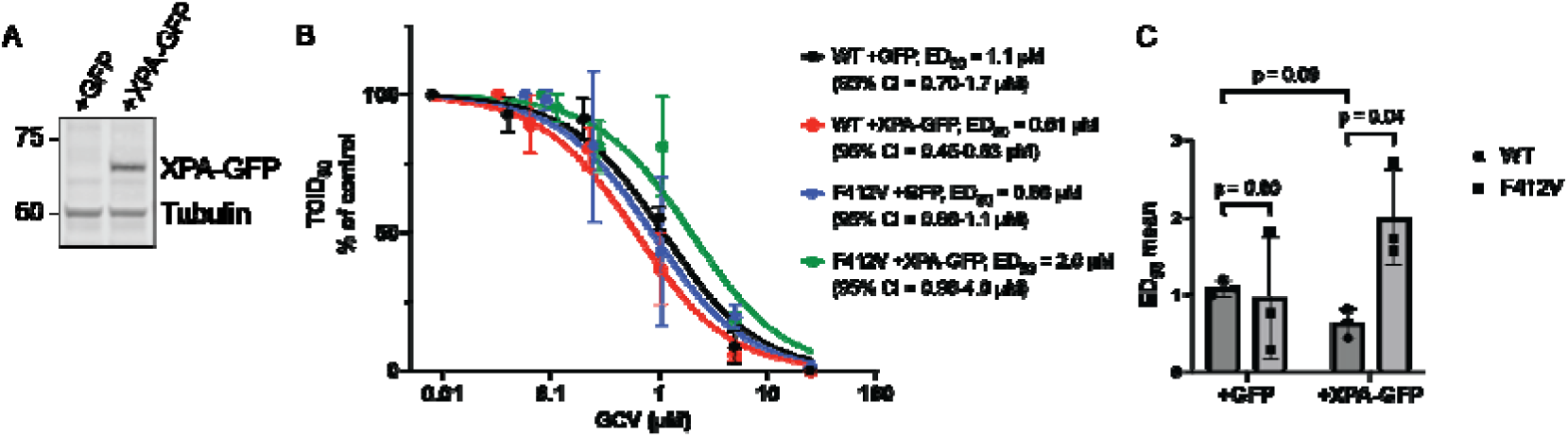
XPA rescue in *XPA*^-^ deficient fibroblasts restores F412V resistance to GCV. (A) Whole cell lysates from *XPA^-^* fibroblasts transduced with GFP-expressing control vector lentivirus (+GFP, left lane) or one expressing XPA-GFP (+XPA-GFP, right lane) were collected to determine XPA expression by immunoblotting. Tubulin serves as a loading control. (B) *XPA^-^* fibroblasts were infected at an MOI of 1 with WT or F412V. At 2 hpi, virus was washed out and media was replaced containing various concentrations of GCV. At four days post infection, cells and supernatant were collected for analysis by TCID_50_. Virus yield at each GCV concentration was compared to the DMSO control for each condition and plotted as percent yield reduction for three biological replicate experiments. Curves were similarly fitted for WT and F412V in each individual experiment to obtain ED_50_ values and CIs for each condition. (C) Those ED_50_ values from three biological replicate experiments were used for a ratio paired t test comparing viruses within each cell type and WT in the two cell types and the Holm-Šídák method was used to correct for multiple comparisons. ED_50_ mean values are plotted with standard deviation error bars.

**Figure 6.**
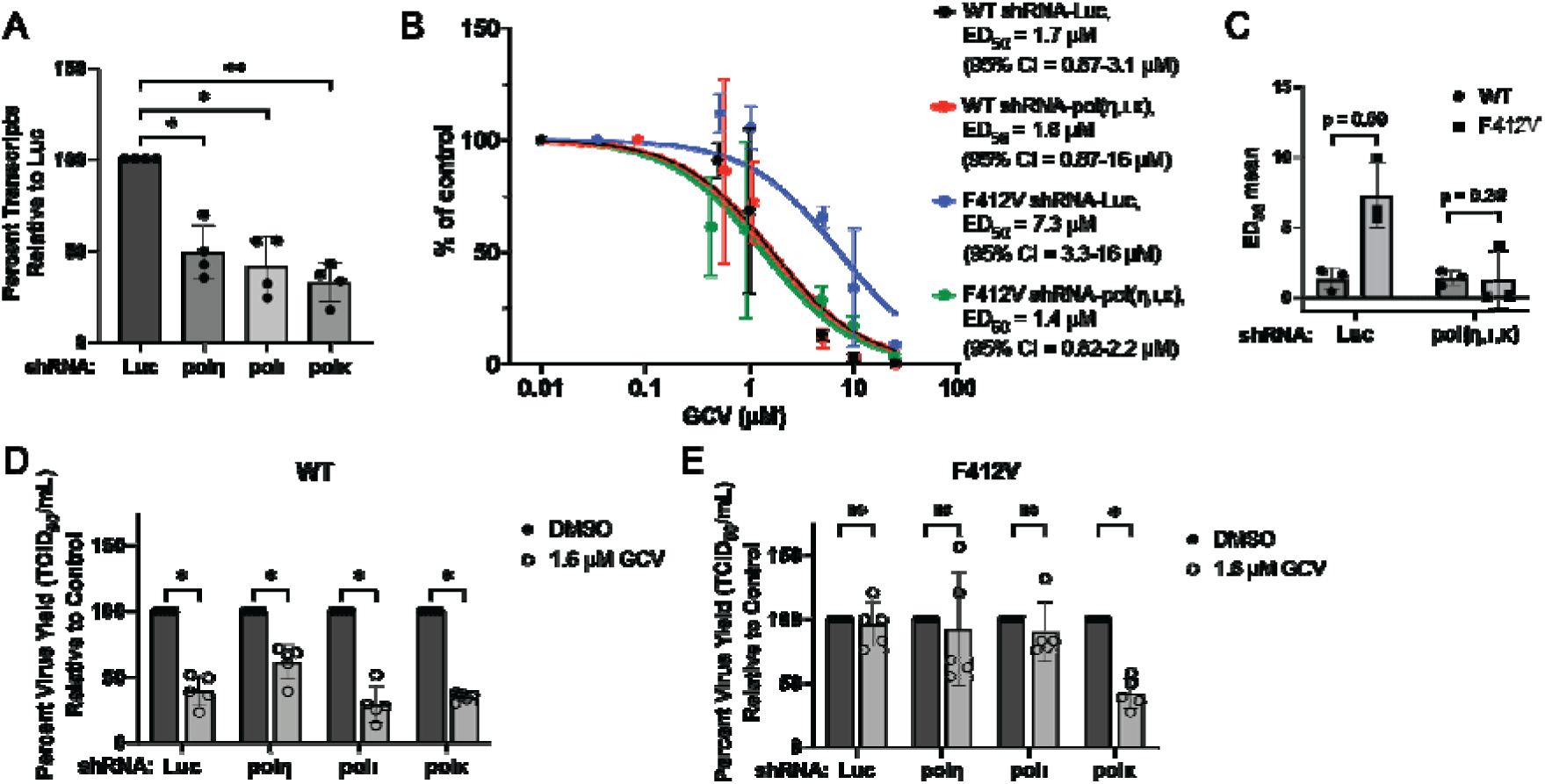
Knockdown of TLS polymerase κ sensitizes Exo-mutant F412V to GCV in fibroblasts. Growth-arrested MRC-5 fibroblasts were transduced with lentiviruses to express shRNAs targeting Luc or three Y-family polymerases: polη, polι, or polκ in combination (A-C) or individually (D and E). (A) Following two days of selection by puromycin (2 µg/mL), cells were lysed and RNA was harvested for analysis by RT-qPCR. Knockdown efficiency was determined by ΔΔCt analysis between Luc and experimental conditions with cellular H6PD as a housekeeping control. Statistical significance was determined by two-tailed, paired *t*-test with Bonferroni correction. Asterisks (* P <0.05, ** P < 0.01) represent statistically significant differences relative to Luc. (B) Transduced fibroblasts were infected at an MOI of 1 with WT or F412V. At 2 hpi, virus was washed out and media was replaced containing various concentrations of GCV. At four days post infection, cells and supernatant were collected for analysis by TCID_50_. For each condition, virus yield at each GCV concentration was compared to the DMSO control and plotted as percent yield reduction for three biological replicate experiments. Curves were similarly fitted for WT and F412V in each individual experiment to obtain ED_50_ values and CIs for each condition. (C) The ED_50_ values from three biological replicate experiments were used for a ratio paired t test comparing viruses within each condition. ED_50_ mean values are plotted with standard deviation error bars. (D-E) Transduced fibroblasts were infected with (D) WT or (E) F412V at an MOI of 1. At 2 hpi, virus was washed out and media was replaced containing either 0.1% DMSO or 0.1% DMSO plus 1.6 µM GCV, and at 4 days post infection, cells and supernatant were collected for analysis by TCID_50_. Percent virus yield in GCV-treated conditions compared to the DMSO control in each condition was plotted for five biological replicate experiments. Statistical significance was determined by a ratio paired t test: *, p<0.05; ns, not significant.

To examine the incorporation of GCV and its anticipated removal, subconfluent HFF cells were serum starved for two days to arrest cells in G0/G1 and then infected at a MOI of 4 with WT, F412V and F412V_R virus (Fig. 2), a protocol that has been shown to greatly reduce synthesis of host cell DNA(44). After 12 h in low serum medium, infected cells were incubated with 0.6 μM [^3^H]-GCV in medium containing 10% FBS, harvesting cells for measurements of incorporated GCV at 48, 72, and 96 h post-infection (hpi). [^3^H]-GCV was washed out at 96 hpi and cells were then maintained in fresh 10% FBS medium lacking GCV. Cells were also harvested at 4h, 6h, and 20h following the removal of radiolabeled GCV and incorporated [^3^H]-GCV was measured (Fig. 2). The amount of [^3^H]-GCV found in DNA is a function of both the rate of incorporation and the rate of removal, and packaging of replicated DNA into capsids would be expected to protect the radiolabel from removal mechanisms. Further, given the requirement for using the relatively high concentrations of GCV that differentiate WT and mutant replication, the specific activity of the [^3^H]-GCV was low in these experiments, so the numbers of cpm incorporated are low. At 48 hpi there was very little incorporation of labeled GCV in cells infected with any of the viruses. Subsequently, in F412V-infected cells, [^3^H]-labeled DNA increased nearly five-fold between 48 and 96 hpi, and the total amount incorporated between 72 and 96 hpi (area under the curve) was significantly greater than that detected from WT- or F412V_R-infected cells (Fig. 2). Notably, during the washout period, the amount of incorporated radiolabel in F412V-infected cell DNA significantly fell (1.4 fold; 29%), while the amount of incorporated radiolabel decreased less (1.2 fold; 17%) for WT or increased for F412V_R, neither of which change was significant (Fig. 2).

One possible explanation for the increased incorporation of GCV into DNA in mutant-infected cells is that the mutant DNA polymerase synthesizes viral DNA more proficiently. To address this possibility, we extracted and sequenced DNA from infected fibroblasts treated with DMSO vehicle without drug and examined the distribution of sequencing reads across the genomes. Read counts were relatively even across the HCMV genome for WT and F412V viruses (Fig. 3A and 3B, top), with the exception of peaks that correspond to regions that include repeat sequences around 50,000 and 240,000 bp on the genome and likely reflect the difficulty of assigning alignment to repetitive sequences, which were uniformly observed in all conditions. Total reads per million were nearly identical not only across the entire genome, but also in the unique sequence region furthest from the origin of replication *oriLyt,* at around 20,000-30,000 bp (Fig. 3C), consistent with there being no defect or advantage in DNA synthesis for the F412V virus.

We also examined the effects of 1.6 μM GCV, which under these conditions usually inhibits WT yield ≥2-fold with little effect on F412V yield (e.g., Fig. 6), on the distribution of sequencing reads for both viruses. In the WT infection treated with GCV, there were significantly fewer reads overall across the genome and especially in the region furthest from *oriLyt* (Fig. 3C; also compare Fig. 3 A and B, top, noting the differences in Y-axes). Strikingly, reads accumulated at ori*Lyt* and decayed rapidly from the *oriLyt* (Fig. 3B), further indicated by the slope of this decay (Fig. 3D), indicative of chain termination. Importantly, in the F412V infection treated with GCV, there was a much less pronounced accumulation of reads at the origin in the presence of GCV and a smaller decrease in total reads and reads distant from *oriLyt*. While the decrease in reads distant from *oriLyt* for the F412V mutant with GCV treatment was statistically significant, there were still significantly more reads than in the WT infection treated with GCV, indicating more complete genome synthesis in the mutant (Fig. 3C, bottom). Moreover, there was a substantially reduced slope of decay from the ori*Lyt* relative to the WT infection treated with GCV (Fig. 3B, D, and E). These results are consistent with not only the F412V DNA polymerase extending DNA chains from GCV-containing primers in vitro (24), but also synthesizing long chains of viral DNA following the incorporation of GCV in infected cells.

Taken together, the results of Figs. 2 and 3 indicate that during F412V infection, the mutant polymerase overcomes chain termination to permit long chain DNA synthesis and increased incorporation of GCV, but that a significant amount of the incorporated drug is subsequently removed from the mutant virus genome, consistent with the possibility that DNA was being repaired following incorporation of GCV.

### GCV resistance of Exo-mutant virus requires XPA

Among major host DNA repair systems, base excision repair (BER), NER, and HR have all been reported to be associated with HCMV infection(11, 12, 45, 46). While certain activities of BER and HR may be downregulated by HCMV to facilitate viral DNA replication and promote genomic variability(45–47), NER is the primary pathway for the repair of a wide range of bulky DNA adducts, including those formed by UV irradiation, which distort the DNA double helix, a property of GCV as well(27, 48). Moreover, an NER-associated factor, XPE also known as DNA damage binding protein 2, is required for efficient HCMV replication(11), and essential components of the NER pathway have been demonstrated to be tightly associated with nuclear viral replication compartments, and to preferentially repair damage to the viral genome over those in host DNA when infected cells are exposed to UV irradiation(12).

Encouraged by these prior studies, we hypothesized that, if NER was responsible for GCV repair, infection of NER-deficient cells would restore the sensitivity of the Exo-mutant virus to GCV. As an initial test of this hypothesis, we used a NER-deficient fibroblast cell line (GM05509, referred to here as *XPA^-^*; Coriell Institute), which was derived from a 31-year-old male patient who carried a mutation in the Xeroderma Pigmentosum group A (*XPA*) gene, and a control fibroblast cell line derived from a healthy, 27-year-old male donor (GM23962, referred to here as Control; Coriell Institute). XPA is an essential protein in the NER pathway, functioning as a scaffold to verify DNA damage and assemble the NER incision complex(49).

We infected Control and *XPA^-^* fibroblasts with WT, F412V, and F412V_R viruses in parallel. Virus replication and sensitivity to GCV was measured using a yield reduction assay, harvesting at 5 dpi to measure infectious progeny yields in the presence of a series of concentrations of GCV. As shown in Fig. 4A in Control cells, F412V was resistant to GCV, while F412V_R and WT exhibited indistinguishable susceptibility to GCV, as expected. In *XPA^-^* cells, the ED_50_’s for all three viruses were noticeably lower. Regardless, resistance of F412V to GCV was largely lost in these cells (Fig. 4B); with the ED_50_ increased ∼3.8-fold in the F412V virus infection relative to WT in Control cells but increased only ∼1.6-fold relative to WT in *XPA^-^*cells (Fig. 4C). While the increased ED_50_ of F412V compared to WT was statistically significant for both control and *XPA^-^*cells, the fold-resistance (ratio of mutant ED_50_ to WT ED_50_) was significantly different between cell types (Fig. 4D). To test the possibility that the change in susceptibility might be caused by slower growth of the mutant virus in *XPA^-^* cells, we measured the replication kinetics of WT and F412V in each cell line in the absence of GCV. Both viruses exhibited similar growth kinetics in the two cell lines during five days of infection, with the mutant virus if anything replicating better in the *XPA*^-^ cells than the normal cells (Fig. S2C).

As the *XPA^-^* and control cells come from different individuals and thus are not isogenic, we then tested whether the loss of XPA protein was responsible for this increased susceptibility of F412V to GCV. We sought to rescue XPA expression by transducing a lentivirus that efficiently expresses XPA fused to green fluorescent protein (GFP) into the *XPA^-^*cell line (Fig. 5A). In yield reduction assays, XPA rescue did not significantly alter GCV sensitivity for the WT virus (Fig. 5B and 5C). However, XPA re-expression restored ganciclovir resistance in F412V infection, increasing the ED_50_ ∼3.2-fold compared to WT infection (Fig. 5C). In the *XPA^-^*cells transduced with a control vector expressing GFP alone, F412V exhibited no significant resistance to GCV, which differs from observations in Figure 4B. We note that for WT infections, there was less than a two-fold difference in virus yields from XPA-transduced ([3.6 ± 0.95] x 10^5^ TCID_50_/mL) and GFP-transduced ([3.0 ± 0.24] x 10^5^ TCID_50_/mL) cells in the absence of GCV, which was not significant (p=0.09). Thus, XPA did not significantly impact replication of the WT virus in this system. Taken together, these data demonstrate that XPA is required for resistance to GCV in Exo-mutant virus infection, suggesting that NER could be responsible for repair of incorporated GCV in the genome of F412V mutant virus.

### Depletion of Y-family polymerase κ restores Exo-mutant susceptibility to GCV

TLS DNA polymerases have enlarged binding pockets and no proofreading function such that they can synthesize through warped DNA landscapes containing adducts, crosslinks, dimers or abasic sites, preventing fork collapse(50–53). We have previously demonstrated a role for host TLS DNA polymerases in viral DNA repair during herpesvirus infection(9). Specifically, we have shown a role for Y-family TLS polymerases—eta (η), iota (ι), and kappa (κ)—in maintaining HCMV genome integrity and introducing single nucleotide variations (SNVs) to viral DNA(9). In addition to the role of Y-family polymerases in TLS, they also contribute to other DNA repair processes, including NER(54).

Therefore, we investigated whether TLS polymerases contribute to resistance to ganciclovir of Exo-mutant HCMV. We performed yield reduction assays on cells transduced with a pool of shRNAs targeting pols η, ι, and κ together or firefly luciferase (Luc) as a non-targeting control. Fibroblasts were growth-arrested prior to transduction to minimize cellular effects due to depletion of polymerases(9). We initially depleted polymerases in combination as they may compete for binding at lesions(55). TLS polymerase transcripts were diminished by at least 2-fold (Fig. 6A), which we previously found to correlate with decreased protein levels(9). Compared to cells transduced with the Luc control, combined depletion of pol(η,ι,κ) increased the susceptibility of the F412V-mutant virus to GCV ∼5-fold, resulting in ED_50_ values similar to those of WT under both conditions. Although this decrease was not statistically significant in our comparison of three biological replicates (Fig. 6B-C), these results are consistent with a role for TLS polymerases in reducing sensitivity of the Exo-mutant HCMV to GCV.

While Y-family pols are recognized for their role in the canonical TLS pathway, they also have roles in other host DNA repair pathways(56). For example, polη can reinitiate DNA synthesis from HR intermediates(57) and polκ can serve as an alternative polymerase to DNA polymerase delta (δ) or epsilon (ε) during the repair synthesis step of NER(54). To understand specific contributions of TLS polymerases, we performed five biological replicates of single-dose yield reduction assays in cells depleted of these polymerases individually at the WT ED_50_ for GCV obtained from Fig. 6B. WT virus exhibited the expected susceptibility to GCV at this dose in cells individually depleted of each TLS polymerase (Fig. 6D). In contrast, although the Exo-mutant virus remained resistant to GCV when polη or polι were individually depleted with yields similar to those seen with the Luc control, GCV reduced F412V virus yield 2.3-fold when polκ was depleted. This reduction was significant, indicating that depletion of polκ restores sensitivity to GCV (Fig. 6E). These results indicate that polκ, but not polη or polι, is required for GCV resistance in Exo-mutant virus infection.

### Polκ depletion also restores Exo-mutant susceptibility to cidofovir

Cidofovir (CDV) can be used as a second-line antiviral drug in cases of GCV resistance. CDV is a nucleotide analog of cytidine monophosphate that, like GCV, inhibits the HCMV Pol through promoting nonobligate chain termination after incorporation of two consecutive molecules into replicating DNA(58). Viruses with Exo-mutant Pol, including a F412V virus, have been reported to be resistant to CDV(42). We sought to understand whether host DNA repair also contributes to CDV resistance. We first established an ED_50_ of 2.5 µM CDV for WT virus in our assays, while the ED_50_ of the F412V mutant was 5.0 µM (Fig. 7A). To determine if polκ has a role in resistance, we performed single dose yield reduction assays (as in Fig. 6D-E) in the presence of ∼ the WT ED_50_ for CDV on cells transduced with shRNAs targeting polκ or Luc. The WT virus remained sensitive to this dose of CDV with either shRNA, as expected (Fig. 7B). However, while F412V was resistant to CDV in the presence of Luc shRNA, F412V virus yields were significantly reduced (2-fold, similar to the WT virus) in CDV-treated cells treated with polκ shRNA (Fig. 7C). Therefore, depletion of polκ restores sensitivity of F412V virus to CDV in addition to GCV. These results suggest at least similar contributions of host DNA repair proteins to both GCV and CDV resistance in Exo-mutant HCMV infection.

**Figure 7.**
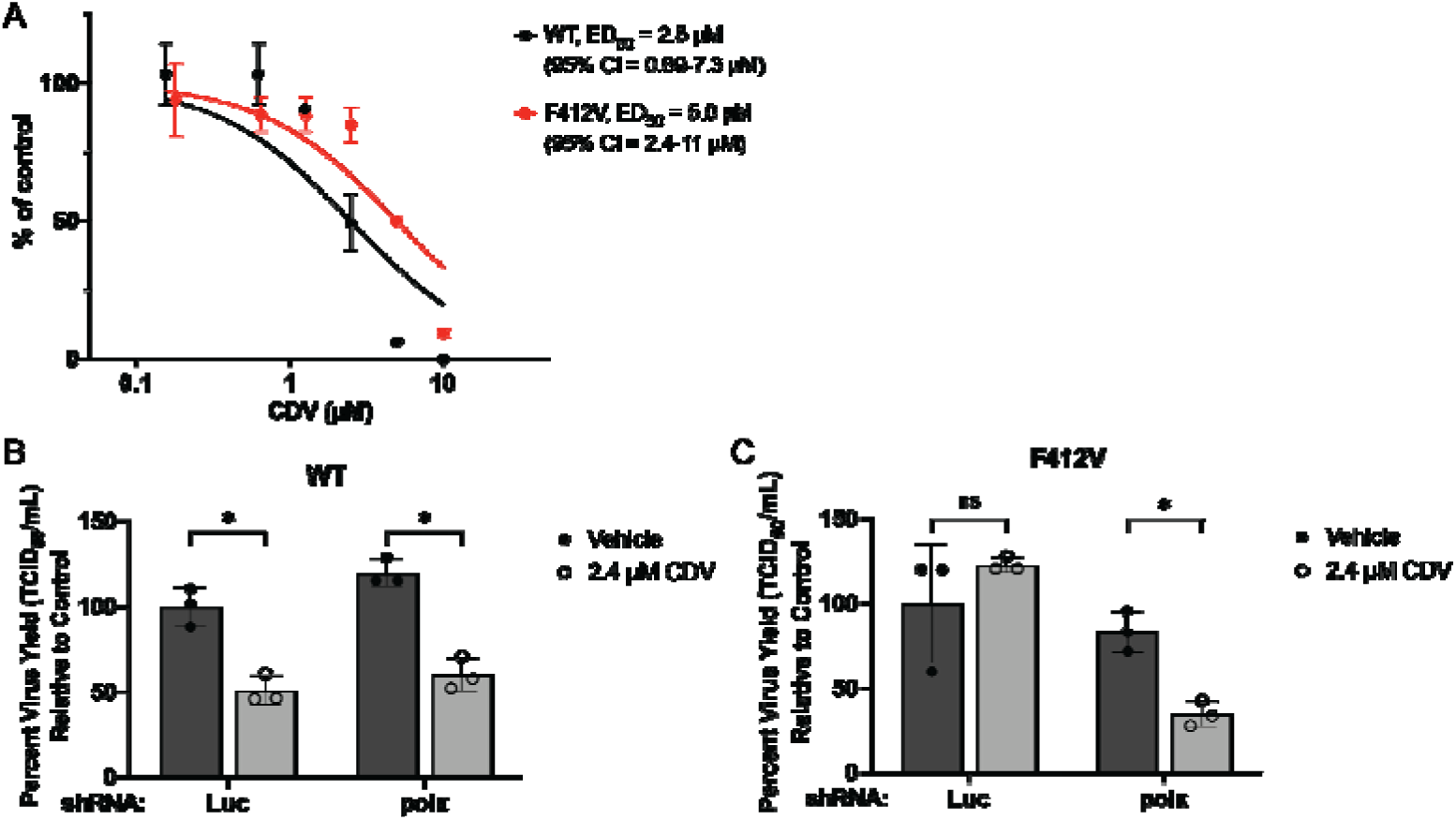
Knockdown of TLS polymerase κ sensitizes Exo-mutant F412V to CDV in fibroblast. (A) MRC-5 fibroblasts were infected at an MOI of 1 with WT or F412V. At 2 hpi, virus was washed out and media was replaced containing various concentrations of CDV. At four days post infection, cells and supernatant were collected for analysis by TCID_50_. For each condition, virus yield at each CDV concentration was compared to the DMSO control and plotted as percent yield reduction for three biological replicate experiments. Curves were similarly fitted for WT and F412V in each individual experiment to obtain ED_50_ values and CIs for each condition. (B-C) Growth-arrested MRC-5 fibroblasts were transduced with lentiviruses to express shRNAs targeting Luc or polκ. These fibroblasts were infected with (B) WT or (C) F412V at an MOI of 1. At 2 hpi, virus was washed out and media was replaced containing either 0.1% DMSO or 0.1% DMSO plus 2.4 µM GCV, and at 4 days post infection, cells and supernatant were collected for analysis by TCID_50_. Percent virus yield in CDV-treated conditions compared to the DMSO control in each condition was plotted for three biological replicate experiments. Statistical significance was determined by a ratio paired t test: *, p<0.05; ns, not significant.

## DISCUSSION

Antivirals are critical in the fight against viral pathogens. Yet, resistance to available antivirals is a growing problem. Mechanisms of antiviral resistance typically involve a mutation in a viral protein, such as the viral polymerase or kinase, that allows the virus to escape antiviral control. However, it is also possible that host processes, including DNA repair, contribute to antiviral resistance. In cancer therapies, DNA repair contributes to drug resistance by removing chain-terminating nucleoside analogs to allow DNA replication and tumor cell proliferation restart(59). However, little is known about the role of host repair mechanisms in the development of antiviral resistance to GCV or other nucleoside analogs. The work presented here in the context of HCMV infection with a GCV-resistant mutant virus serves as a new model to explore this underappreciated relationship. Here, we demonstrate that when HCMV acquires resistance to GCV through a mutation that compromises viral DNA polymerase exonuclease activity, two host proteins, XPA and polκ, are necessary for this resistance. This finding implies that neither protein is sufficient for resistance and that at least one host DNA repair pathway, NER, in which both XPA and polκ participate, and possibly a second, TLS, mediate this resistance. Building on this finding, we show that polκ also mediates Exo-mutant resistance to CDV, suggesting a broader role for host proteins in viral DNA repair and antiviral resistance.

The NER pathway is crucial for excising bulky DNA adducts, including cyclopyrimidine dimers, which kink the DNA double helix(48). Because GCV is toxic to DNA synthesis and its incorporation can also distort the double helix(27), NER was a top candidate among host repair pathways that could excise GCV incorporated into DNA and permit subsequent rounds of viral replication. Indeed, we found that the Exo-mutant virus, F412V, exhibits reduced resistance to GCV, with an ED_50_ similar to that of WT virus, in cells deficient in NER due to inactivation of the critical *XPA* gene. The NER pathway is active during HCMV replication in fibroblasts and sufficient to repair cyclopyrimidine dimers in viral DNA induced by UV treatment(11, 12). In support of this, O’Dowd and colleagues also demonstrated that various NER proteins associate with viral nuclear replication compartments (12). Moreover, DDB2, an important component of the Cul4 ubiquitin ligase complex with roles in the NER pathway, is important for efficient viral DNA synthesis(13). These findings are consistent with the role for NER in GCV-resistance defined here.

Our results together with the previous studies suggest a model (Fig. 8) where the NER apparatus assembles at locations of internally incorporated GCV and excises the nucleoside analog, leaving a gap that is then filled (Fig. 8C). There are two subpathways for this mode of repair: global genome NER, which occurs anywhere in the genome, and transcription-coupled NER, which occurs on actively transcribed genes(48). It remains to be determined whether global genome NER or transcription-coupled NER is more dominant in the context of Exo-mutant infection.

**Figure 8.**
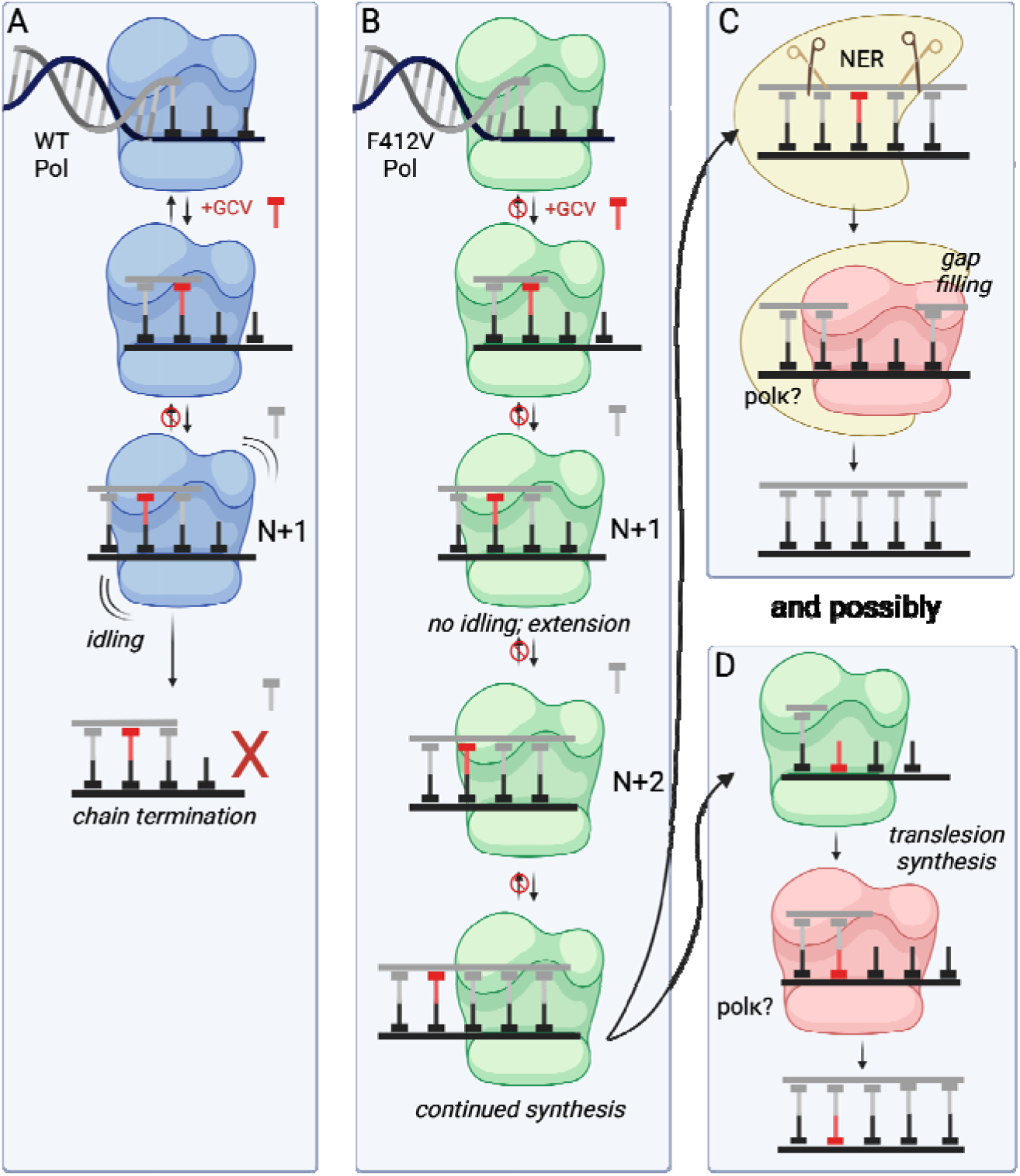
Model for host-mediated repair of GCV-containing viral genomes. During HCMV replication in the presence of GCV (A) the WT Pol incorporates GCV and an additional nucleotide (N+1). Synthesis terminates at the N+1 position due to idling (addition of the next (N+2) nucleotide followed by its rapid excision) that depends on the WT Pol exonuclease activity. (B) An exonuclease-deficient Pol, such as that of the F412V mutant virus, also incorporates ganciclovir during DNA synthesis. However, following incorporation of the N+1 nucleotide, the F412V Pol does not idle and, instead, continues synthesis. While these GCV-containing templates should not be competent templates for subsequent rounds of synthesis (and possibly transcription), the corresponding resistance mechanism must allow these processes. We show that resistance requires specific host mechanisms of DNA repair. From this work, we propose that (C) the host repair mechanism, NER, can excise GCV-containing DNA from the newly synthesized strand (gray) with polκ completing the gap filling step. (D) Additionally, if a template strand (black) contains GCV in a subsequent round of DNA synthesis, we propose that polκ might complete replication across from the GCV lesion through its activity in translesion synthesis.

It is not clear how NER or any other DNA repair pathway might impact GCV efficacy in HCMV infection expressing WT Pol. While susceptibility of WT virus to GCV was ∼2-fold lower in Control cells relative to *XPA^-^*cells (Fig. 4), the XPA- and XPA+ cells are not isogenic. We did not observe a significant difference in WT GCV susceptibility between the *XPA^-^* cells transduced with XPA or control (Fig. 5). Therefore, it is more likely that NER has little, if any, role in removing GCV from genomes where efficient chain termination occurs. Further, there was no significant loss of radiolabeled GCV from DNA in WT- and revertant-infected cells following washout, although it is possible that poor incorporation of GCV by these viruses during the labeling period made it difficult to detect any repair. Either way, the circumstances for repair following incorporation of GCV by WT Pol are rather different than those for Exo-mutant Pol as in the former case the enzyme idles at the N+1 position with the GCV just one base removed from the primer-terminus in the active site. In the latter case the polymerase and replication fork have traveled away from the inserted GCV (Fig. 8C), allowing room for access to and triggering of NER. Moreover, due to chain termination, the WT Pol would create stalled replication forks, at least following incorporation of GCV into the leading strand, and resolution and repair of those stalled forks are likely to be even more complicated and slower than NER.

Our study further demonstrates a role for TLS polymerases, particularly polκ, in HCMV resistance to GCV and CDV treatment. A parsimonious model for this role would be that polκ acts during NER as the polymerase for the nucleotide insertion step for gap filling following removal of the lesion (Fig. 8C), and might be particularly important in the G1/G0 phase of the cell cycle (54, 60), where HCMV initiates its replicative program prior to blocking full entry into S-phase(61, 62). Moreover, recruitment of polκ during NER requires monoubiquitinated PCNA, which we previously found to be induced during replicative infection, particularly at the initiation of viral DNA synthesis(63). This provides further support for repair of GCV-containing HCMV DNA by the NER pathway. Interestingly, polη, which is encoded by the Xeroderma pigmentosum gene *XPV*, can serve as a TLS across from lesions, including those repaired by NER(64). However, its knockdown alone did not restore sensitivity to the Exo-mutant virus (Fig. 6E). This seemingly negative result is consistent with NER being more important than bypass synthesis by TLS.

However, we note that in many other contexts and possibly following the incorporation of GCV, DNA damage is bypassed by TLS Y-family polymerases, which lack 3’-5’ exonuclease (proofreading) activity and synthesize through bulky lesions and adducts in the template strand where high-fidelity polymerases would otherwise stall(50, 53). Therefore, TLS could contribute to resistance in a subsequent round of DNA replication by facilitating synthesis across the GCV-containing template strand during replication (Fig. 8D), which we have found is performed inefficiently at best by HCMV Pol. Importantly, in addition to lacking proofreading activity, Y-family polymerases have enlarged binding pockets, causing them to be error-prone. In a previous study, we observed decreased single nucleotide variants (SNVs) in viral DNA when pols η, ι, and κ were depleted, which suggests that their error-prone function may contribute to viral genome diversity(9). In the context of Exo-mutant virus infection, if TLS pols function in DNA synthesis opposite GCV-containing template DNA, then increased SNVs would be expected on the newly synthesized viral DNA. Regardless, TLS does not explain the removal of GCV from viral DNA that we observed, which we attribute at least in part to NER. Defining specific contributions of polκ and various host repair pathways will be important in future work, with emphasis on determining the relative roles of NER and TLS and their effects on GCV incorporation and removal in both viral and host DNA, as well as defining mechanisms of Y-family pol functions on viral DNA to understand how they modulate viral genome integrity, especially in the presence of GCV.

We note some possible limitations to our study. Our biochemical assays used only a single primer-template to test the ability of Pol synthesize across from GCV in the template, so although our DNA sequencing studies and the requirement for host DNA repair proteins for the resistance of F412V virus suggests that the inability of HCMV Pols to synthesize across from GCV occurs often in the viral genome, we cannot rule out effects of sequence context on this phenotype. We tested only a single Exo-mutant, so that although it seems highly likely that the GCV-resistance of other Exo-mutants or the A987G mutant that overcomes idling by more rapid polymerization(25) would also depend on XPA and polκ, we cannot say that for certain. We also only used fibroblasts for these studies, so it is possible that resistance may not depend on XPA and polκ or host repair at all in different cell types, although we do note that Exo-mutants do cause treatment-refractory disease in many tissues.

The use of GCV with HSV-TK as a cancer therapy provides an important foundation for our study. In investigations of this anticancer treatment, multiple groups discovered that intrinsic differences in the DNA repair landscape of various tumors contribute to variable susceptibility to this form of suicide gene therapy. For example, Tomicic et al. reported that polymerase beta (β), which has key roles in the BER pathway, contributed to resistance against GCV-induced cytotoxicity in various cell culture models(30). In line with this, chemical inhibition or knockout of polβ sensitized cells to GCV and HSV-TK treatment(30). Interestingly, HCMV downregulates specific aspects of the BER pathway(45) while inducing or hijacking NER(12) (and this study). Taken together, these findings underscore the importance of excision pathways in removing incorporated GCV, and possibly other nucleoside analogs, as a mechanism of resistance not only for nucleoside analogs that treat viral diseases, but also cancer.

Targeting DNA repair pathways is a common strategy for anticancer therapy, especially for tumors which have become resistant to chemotherapies. Various studies have specifically highlighted NER(65) and TLS(55, 66) as potential therapeutic targets that could synergize with genotoxic chemotherapies. Our findings shed light on the importance of these pathways in antiviral resistance and raise the possibility that small molecule inhibitors targeting these host DNA repair pathways might improve GCV or CDV therapy for patients who have acquired resistance due to exonuclease-deficient HCMV. Potential barriers to development of such inhibitors for this indication include the availability of other antiviral drugs that can treat such mutant viruses, the requirement for testing to determine the presence of this mechanism of resistance, and the risks of unrepaired host DNA damage in immunologically impaired populations. Regardless, further exploration of this possibility seems warranted.

## MATERIALS AND METHODS

### Protein purification

WT HCMV UL54 pol and mutant pols (F412V, L545S and A987G) were expressed as GST tagged proteins and purified using affinity chromatography as described previously(24, 25).

### Oligonucleotides

Primer-template T1 was purchased from integrated DNA technologies and primer-template T5 was synthesized by ChemGenes using GCV phosphoramidite prepared as described by Marshalko et al. (67).

### Polymerase assay

Polymerase assays for testing chain extension on T1 and T5 by HCMV WT and mutant polymerases used the methods and conditions described previously(24, 25) with some modifications. Briefly, the reactions were performed in 5 μL volumes and contained 1-2 pmol of the indicated primer-templates, 5’ end-labeled using [γ - ^32^P]-ATP (PerkinElmer) and T4 polynucleotide kinase (New England Biolabs), 500 fmol of each Pol with a twofold molar excess of UL44ΔC290 (kindly provided by Gloria Komazin-Meredith, Harvard Medical School), 250 μM each of dATP, dCTP, dGTP and dTTP, 50 mM Tris (pH 8.0), 1mM DTT, 100 mM KCl and 40 μg/mL BSA. To test chain extension with exonuclease-deficient T7 DNA polymerase (kindly provided by Charles Richardson, Harvard Medical School), the reactions were performed in a 10 μL volume containing 4 pmol of the indicated radiolabeled primer-template, 1 pmol T7 DNA Pol, 250 μM each of dATP, dCTP, dGTP and dTTP, 40 mM Hepes (pH 7.6), 5 mM DTT, 100 mM NaCl. For all assays, reactions were initiated by adding 10 mM MgCl_2_ and, after incubation at 37° C for 10 min, quenched using 5 μL of stopping buffer (assays with HCMV Pols) or after 2 min, using 10 μL stopping buffer (assays with T7 DNA polymerase) as described previously(24, 25). The products in stopped reactions were resolved on a 20% denaturing polyacrylamide gel and the gel images were acquired using an Amersham Typhoon 5 biomolecular imager (GE Healthcare). Densities of bands in Fig.1 were quantified using ImageJ. The incorporation efficiencies for T7 Pol, WT HCMV Pol and mutants on T1 or T5 were calculated by dividing the density of the full-length product by that of the corresponding primer-template in each lane.

### Cells and viruses

Human foreskin fibroblast (HFF) cells (Hs27, American Type Culture Collection [ATCC]) were propagated in DMEM containing 10% fetal bovine serum (FBS), 100 U/mL penicillin, and 100 µg/mL streptomycin. Primary human lung MRC-5 fibroblasts (ATCC CCL-171) were cultured in DMEM supplemented with 10% FBS, 10 mM HEPES, 1 mM sodium pyruvate, 2 mM L-alanyl-glutamine, 0.1 mM nonessential amino acids, 100 U/mL penicillin, and 100 µg/mL streptomycin. Control and *XPA^-^* fibroblast cell lines, GM05509 and GM23962 (Coriell Institute) were maintained following the guidance from the vendor. HCMV 53-F BADGFP(68) served as the WT virus. The virus carrying the *UL54* mutation F412V was constructed by introducing the mutations into the bacmid 53F pBADGFP(68) with the primers listed in Table S1 followed by two-step red recombination(69, 70). To generate the rescued virus F412V_R, wild type sequences were restored to the mutant BAC using the PCR primers listed in Table S1 and the same methods cited above. Each virus was titrated as described previously(71).

### Antiviral activity and replication kinetics

For Figures 4 and S1A, the antiviral activity of GCV against WT HCMV, F412V and F412V_R was assessed using yield reduction assays as described previously(72) with modifications as follows: 1x10^5^ HFF (Fig. S1A), XPA deficient cells (GM05509) or Control cells (GM23962) (Fig. 4) in each well of a 24-well plate were infected with virus at MOI 2. After 2h incubation at 37° C, the virus was removed and the cells were incubated with 1mL fresh DMEM medium containing either 1% DMSO or various concentration of GCV (Fisher Scientific) and 1% DMSO. At 5 dpi, the supernatants were harvested and the titers of virus were determined using standard plaque assays. The ED_50_ for GCV against each virus was calculated by normalizing the number of plaques at each concentration of GCV to those in DMSO control using nonlinear regression in GraphPad Prism 9.5 for MacOS.

The replication kinetics of these viruses with either no drug, 0.3 μM, or 0.6 μM GCV were assessed as just described with no drug only using either HFFs (Fig. S2A and S2B) or with Control or *XPA^-^* cells (Fig. S2C) except that supernatants were harvested at 24, 48, 72, and 96 hpi and the determined titers were plotted against time.

For Figure 5, to establish the XPA rescue and control cell lines, GM05509 fibroblasts (*XPA*^-^) were transduced in a 10 cm dish at MOI 3 with lentiviral particles to express XPA-GFP (Origene RC204872L2V) or unfused GFP control (Origene PS100093V) and 10 µg/mL polybrene. Forty-eight hours following transduction, growth medium was replaced with medium containing 2 µg/mL puromycin. Forty-eight hours later, surviving cells were collected and maintained in growth medium containing puromycin. These cells were seeded into 12-well plates at 200,000 cells per well and then infected with HCMV at an MOI of 1 the next day. Following two hours of infection, media was replaced with media containing GCV (0.008-25 µM) in 0.1% DMSO or a 0.1% DMSO vehicle control. At four days post infection, cells and supernatant were harvested for analysis of virus yield by 50% Tissue Culture Infectious Dose (TCID_50_) assay. Briefly, 10,000 cells were seeded per well into a 96-well plate. The next day infected cell lysates were prepared in 10-fold dilutions (8 total, one per row) and then overlaid onto the cells. At fourteen days post infection, plates were scored by assessing the number of GFP-positive wells. Virus yields were calculated using the Reed-Muench method(73). The ED_50_ for each virus against GCV was calculated by normalizing the virus yield at each GCV concentration to that of the respective DMSO control and plotted using GraphPad Prism 10 for MacOS.

For Figures 6 and 7, MRC-5 fibroblasts were seeded into 12-well plates at 200,000 cells per well. Cells were transduced with lentiviral particles delivering shRNA constructs targeting luciferase or polη, polι, and/or polκ as previously described(9). At four days post-transduction, cells were infected or mock-infected with HCMV at an MOI of 1. Following two hours of infection, media was replaced, and cells were treated with GCV (0.008-25 µM) in 0.1% DMSO or a 0.1% DMSO vehicle control (Fig. 6); or cells were treated with CDV (0.15-10 µM) in water or a water vehicle control (Fig. 7). At four days post infection, cells and supernatant were harvested for analysis of virus yield by TCID_50_ assay. The ED_50_ for each virus against GCV was calculated by normalizing the virus yield at each GCV concentration to that of the respective vehicle control and plotted using GraphPad Prism 10 for MacOS. For single dose assays in Fig. 6D-E and 7B-C, a similar infection approach was implemented, but cells were treated with a single dose of GCV (1.6 µM) or CDV (2.4 µM) or respective vehicle control. Virus yield at four days post infection was compared between the two treatment groups.

### Assays for incorporated GCV in infected HFF cells

5x10^5^ HFF cells in each well of a six-well plate were incubated in DMEM with 0.2% FBS for 48h. After infection with WT, F412V and F412V_R at an MOI of 4, virus was removed after two hours, and cells were overlaid with DMEM with 0.2% FBS and incubated at 37° C. At 12 hpi, the medium was replaced with DMEM containing 0.6 μM [^3^H]-GCV (Moravek Inc.) and 10% FBS and cells were incubated until 96 hpi; then rinsed with PBS and incubated with fresh DMEM medium supplemented with 10% FBS without drug for 20h to wash out the GCV. At 48, 72, and 96 hpi and then 4h, 6h and 20h following removal of GCV, cells were harvested by trypsinization, and treated with 0.4 N perchloric acid at 4° C overnight to precipitate macromolecules including DNA. Following centrifugation at 15,000 xg the acid-insoluble pellets were suspended in cold water, 1N NaOH was added to neutralize and solubilize the pellets, and the amount of radiolabel per well of cells was measured using a scintillation counter. The kinetics of incorporation and loss were analyzed and graphed using GraphPad Prism 9.5 for MacOS.

### Genomic sequencing and computational analysis

MRC-5 fibroblasts were seeded onto 15 cm dishes and infected with HCMV (WT or F412V) at an MOI of 1. Following two hours of infection, the inoculum-containing medium was replaced with fresh medium containing 1.6 µM GCV in 0.1% DMSO or a 0.1% DMSO vehicle control. At four days post infection, cells were washed with PBS and collected in DNA lysis buffer containing 200 μg/ml proteinase K by manual scraping. After a 2 h, 55°C incubation for proteinase K digestion, cellular and viral DNA were isolated using phenol–chloroform extraction.

Each sample was sequenced on a NextSeq 2000 Illumina short read sequencer to yield paired-end short reads. The sequencing reads were aligned to the human reference genome GRCh38 using Bowtie2 v2.5.1 with the following parameters: --very-sensitive for sensitive alignment, --seed 1 for seeding alignment. After alignment, reads aligned to the human genome were filtered out using Samtools v1.17. The following parameters were used: -f UNMAP,MUNMAP to extract reads that were not aligned to the human genome, and -bh to output the filtered alignments in BAM format.

Total coverage at each genomic position was calculated by summing coverage across all strands and mapping categories. The genome was segmented into non-overlapping 1000bp bins. For each bin, mean total coverage was computed. Mean coverage per bin was normalized by the median coverage of the entire sample to control for differences in sequencing depth, allowing comparison across samples. The distance of each bin’s midpoint from either side of the origin of lytic replication (OriLyt; positioned at 140,000bp) was computed as an absolute value. For each sample, the log-transformed normalized coverage was modeled as a function of distance from OriLyt via linear regression, where the slope quantifies the rate of coverage decay with increasing distance from *OriLyt*. Nonparametric LOESS smoothing was performed to visualize coverage trends as a continuous function of distance separately for each group. The slopes estimated per sample were compared across groups using pairwise permutation testing (with BH correction) to assess significant differences in replication bias patterns between conditions.

### Immunoblotting

Whole cell lysates were extracted using radioimmunoprecipitation assay (RIPA) lysis buffer (Pierce® formulation) and manual scraping. 50 µg of lysate in SDS sample buffer was loaded onto precast 4-12% bis-tris gels (ExpressPlus™; GenScript) and then proteins were separated by electrophoresis and transferred onto 0.45 µm pore size PVDF membranes (Immobilon®-FL; Millipore). Proteins of interest were detected using monoclonal primary antibodies: XPA (rabbit mAb, Cell Signaling Technology #14607; used at 1:1000) and tubulin (mouse mAb clone DM1A, Sigma #T9026; used at 1:2000) were used. Mouse DyLight™ 680 (Invitrogen #35519) and rabbit DyLight™ 800 (Invitrogen #SA5-10036) conjugated secondary antibodies were used. Images were obtained using a LI-COR Odyssey CLx scanner and analyzed using ImageStudio software.

### RT-qPCR

MRC-5 fibroblasts were seeded into a 12-well plate at 200,000 cells per well and then transduced with lentiviral particles delivering shRNA constructs as described above. At four days post transduction, cells were washed in PBS and then lysed in 400 µL of DNA/RNA Lysis Buffer from Zymo Research. DNA and RNA were isolated according to manufacturer instructions for ZR Duet DNA/RNA Mini Prep Kit (Zymo Research). cDNA was synthesized from RNA using Transcriptor First Strand cDNA Synthesis Kit (Roche) and quantified using the ΔΔCt method relative to the cellular housekeeping control, H6PD.

## ACKNOWLEDGMENTS

This work was supported by funding from the National Institutes of Allergy and Infectious Diseases, including R01 AI019838 to D.M.C., R01 AI177392 to F.G. and G.B., R37 AI079059, AI079059-14S1, and R01 AI177392002S1 to FG, and R03 AI140048 to H.C. This work was also supported by a Munck-Pfefferkorn grant from the Geisel School of Medicine at Dartmouth. We are grateful for technical assistance from Seamus McCarron, Suresh Srivastava of ChemGenes Corporation, Wilmington, MA, for expertise in the synthesis of ganciclovir-containing oligonucleotides, Charles Richardson for provision of exonuclease-deficient T7 DNA polymerase, and invaluable encouragement from the late G. Peter Beardsley.

