## Supplementary Data for "Host DNA repair factors empower a mechanism of antiviral nucleoside analog resistance"

### Supplemental Information

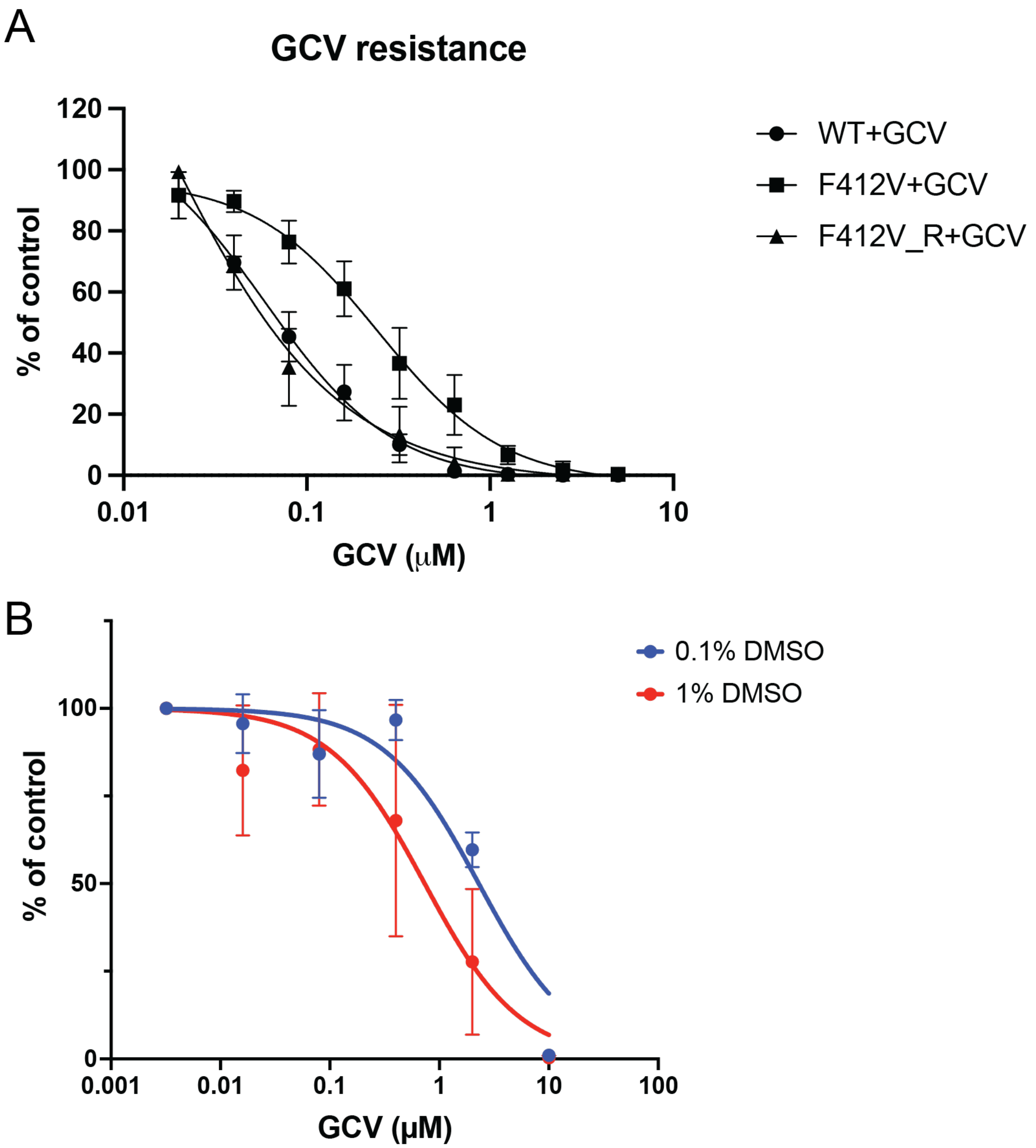


**Figure S1. Sensitivity of WT and F412V to GCV and DMSO.** (A) HFF cells infected with WT, F412V and F412V_R viruses were incubated with GCV in 1% DMSO-containing medium at various concentrations for 5 days. Then the supernatants were harvested and viral titers were quantified using standard plaque assay. Curves were fit using nonlinear regression and $ED_{50}$ values were calculated using Prism 9 for MacOS. Error bars represent standard deviation for triplicates. The $ED_{50}$ value for F412V was ~5-fold higher than those of the other two viruses. (B)

MRC-5 fibroblasts were infected with WT HCMV at an MOI of 1. At 2 hpi, virus was washed out and media was replaced containing various concentrations of GCV in 1% DMSO or 0.1% DMSO-containing medium. At four days post infection, cells and supernatant were collected for analysis by $TCID_{50}$. For each condition, virus yield at each GCV concentration was compared to the no GCV control and plotted as percent yield reduction for three biological replicate experiments.

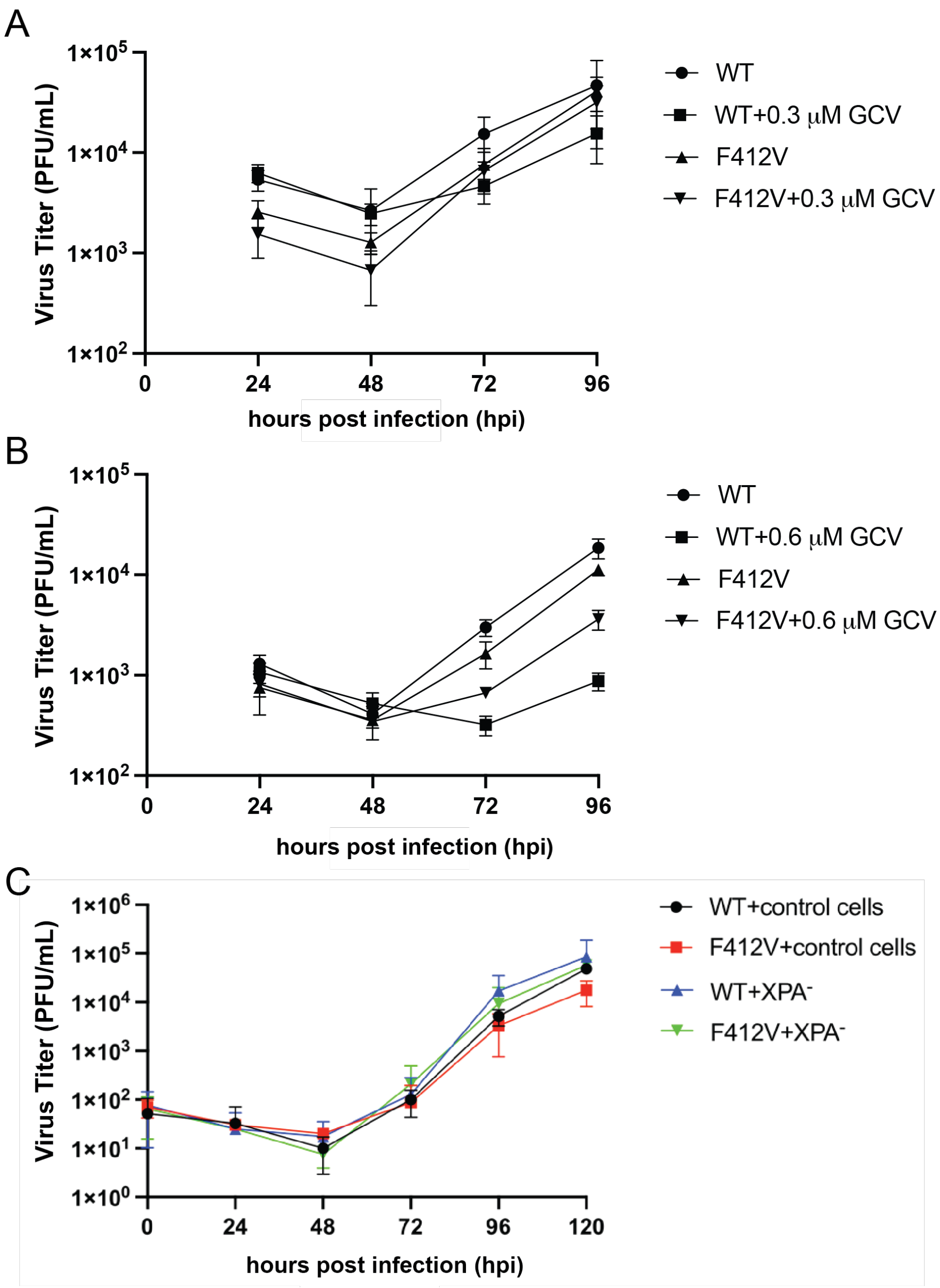


**Figure S2. Viral growth kinetics with and without GCV treatment.** Either HFFs (A and B) or Control and *XPA*⁻ cells (C) were infected with WT or F412V at an MOI of 2 and incubated in the presence of either the absence of ganciclovir or the concentrations of ganciclovir shown in 1% DMSO-containing medium (A and B) or only the absence of ganciclovir (C), harvesting supernatants at the indicated hpi, followed by titration using plaque assays. Error bars represent standard deviations for triplicates.

| Construct | Forward primers | Reverse primers |
|---|---|---|
| BAC_UL54-F412V | 5’TTTCAACGGTACGCGCCGGCCTTTGTGACCGGTTACAACATCAACTCTGTTGACTTGAAGTAGGGATAACAGGGTAATCGATTT 3’ | 5’GTCCACCTTATACAGGTACTCGAGACGCGTGAGGATGTACTTCAAGTCAACAGAGTTGATGCCAGTGTTACAACCAATTAACC 3’ |
| BAC_UL54-V412F | 5’TTTCAACGGTACGCGCCGGCCTTTGTGACCGGTTACAACATCAACTCTTTTGACTTGAAGTAGGGATAACAGGGTAATCGATTT 3’ | 5’GTCCACCTTATACAGGTACTCGAGACGCGTGAGGATGTACTTCAAGTCAAAAGAGTTGATGCCAGTGTTACAACCAATTAACC 3’ |

**Table S1. Primers used for BAC recombination.**
